# A mechanical model of the half-sarcomere which includes the contribution of titin

**DOI:** 10.1101/435420

**Authors:** Pertici, M. Caremani, M. Reconditi

## Abstract

The evidence, in both resting and active muscle, for the presence of an I-band spring element like titin that anchors the Z line to the end of the thick filament did not yet produce a proper theoretical treatment in a complete model of the half-sarcomere. The textbook model developed by A.F. Huxley and his collaborators in 1981, which provides that the half-sarcomere compliance is due to the contribution of the compliances of the thin and thick filaments and actin-attached myosin motors, predicts that at any sarcomere length the absence of attached motors results in an infinite half-sarcomere compliance, in contrast with the observations. Growing evidence for the presence of a titin-like I-band spring urges the 1981 model to be implemented to include the contribution of this element in the mechanical model of the half-sarcomere. The model described here represents a tool for the interpretation of measurements of half-sarcomere compliance at long sarcomere lengths and for investigations of the possible role of titin as the mechano-sensor in thick filament regulation.

## INTRODUCTION

In the striated (skeletal and cardiac) muscle, the contractile proteins myosin and actin are organized respectively in well-ordered thick and thin filaments in the sarcomere, the ca 2 μm long structural unit of muscle in which bipolar arrays of myosin II motors emerging from the thick filaments overlap with the thin filaments originating from the Z line bounding the sarcomere. Force and shortening during muscle contraction are due to cyclical ATP-driven interactions of the globular portion (the head) of the motors with the nearby actin monomers in the thin filaments. In each half-sarcomere (hs), the myosin motors are mechanically coupled as parallel force generators via their attachment to the thick filament, constituting, together with the interdigitating thin filaments and the other cytoskeleton and regulatory proteins, the basic functional unit of muscle (Fig. 1). Stiffness measurements at the level of the half-sarcomere directly inform on the stiffness and the number of actin-attached motors (cross-bridges) under the condition that they are the only source of compliance in the half-sarcomere (Huxley and Simmons 1971; Ford et al. 1977). In contrast to this idea, X-ray diffraction experiments indicated that thin and thick filaments under stress (Huxley et al. 1994; Wakabayashi et al. 1994; Reconditi et al. 2004; Huxley et al. 2006; Piazzesi et al. 2007) extend by 0.23–0.26% for a force change equivalent to *T*_0_, the maximal force developed in an isometric tetanic contraction. Under these conditions the contribution of actin-attached cross-bridges and myofilaments to half-sarcomere compliance (*C*_hs_) can be defined, in the framework of the model developed by Ford and colleagues (Ford et al. 1981; FHS1981 hereafter), in which, assuming that the compliance of the array of cross-bridges is not smaller than the cumulative compliance of the actin and myosin filaments, *C*_hs_ can be calculated by the sum of the equivalent compliances of the three elements (see Methods). The simple useful formulation of this equation is:

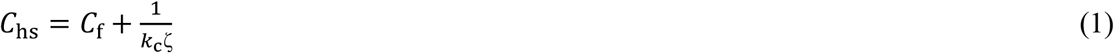

**Fig. 1.**
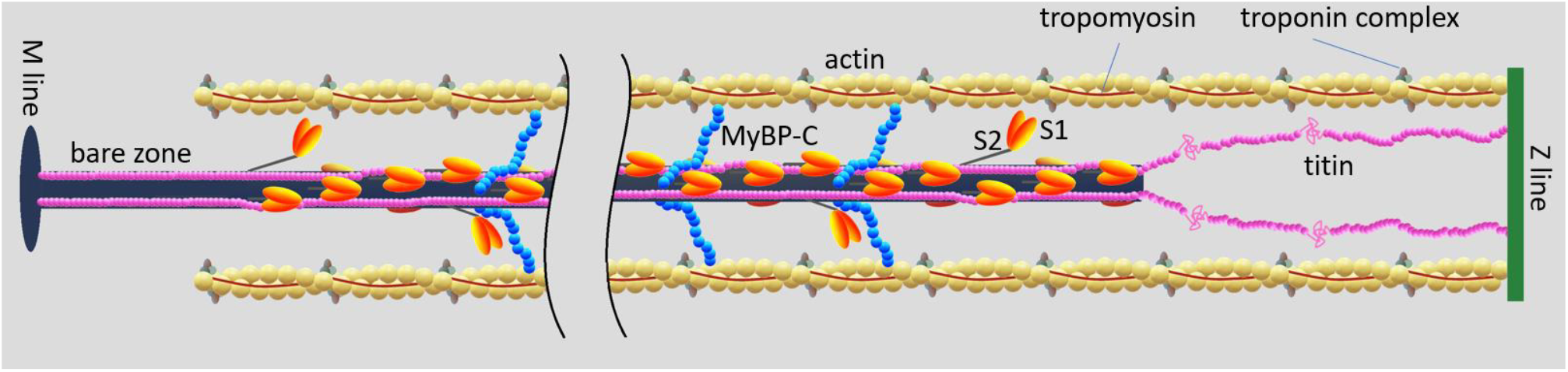
Schematic representation of the half-sarcomere protein assembly. On the thin filament (yellow, formed by the actin monomer polymerization in a double helix with a period of 73 nm) are shown the regulatory proteins tropomyosin (red) and troponin complex (light and dark gray and brown). On the thick filament (dark blue) the S1 globular head domains of myosin (orange) emerge with a three stranded helical symmetry with 43 nm periodicity. The Myosin Binding Protein C (MyBP-C, blue) lies on the proximal 1/3 of the thick filament with the C-terminus and extends to thin filament with the N-terminus. Titin (pink) in the I-band connects the Z line at the end of the sarcomere (green) to the tip of the thick filament and in the A-band runs on the surface of the thick filament up to the M-line at the centre of the sarcomere. To see this figure in color, go online.

Where *C*_f_ is the equivalent filament compliance, *k*_c_ is the stiffness per unit length of the array of the cross-bridges and ζ is the length of overlap of thin filament with the cross-bridges array in each half-sarcomere.

More recently, hs stiffness measurements by means of fast length oscillations applied to single muscle fibers during the development of an isometric tetanus have shown that at low forces, when the number of attached motors is relatively low, a significant contribution to *C*_hs_ emerges from another elastic element the compliance of which, *C*_p_, is functionally in parallel with that of the attached motors (Colombini et al. 2010; Fusi et al. 2010, 2014). The value of *C*_p_ was somewhat controversial depending on the protocols used to estimate it. When stiffness measurements were made at the level of a selected population of sarcomeres in an isometrically contracting single frog fibre at 4°C (plateau force *T*_0_ ~150 kPa), it resulted to be 200-300 nm/MPa (11,12), that is ~20 times larger than the compliance of the array of motors attached at *T*_0_ (11.5 nm/MPa). Such a relatively large compliance of the parallel elasticity explains why its contribution emerges only at low forces when the stiffness of the motor array becomes comparable to that of the parallel element. To consider the contribution of this element to the *C*_hs_ requires only a slight modification of the FHS1981 model. Namely, *C*_hs_ can be interpreted as the series of the filament compliance, *C*_f_, and the compliance resulting from the parallel arrangement of the force-generating cross-bridges and the new element:

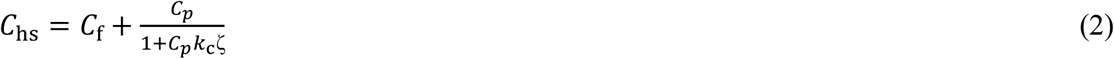

This element could be associated to the presence of links connecting the thin and thick filaments in the A-band, like either a fraction of weakly bound, no-force generating motors (Colombini et al. 20101; Fusi et al. 2017), or the thick filament accessory protein myosin-binding protein C (Fig 1), which has shown to undergo dynamic interactions with the thin filament (Offer et al. 1973; Moos et al. 1981; Yamamoto 1986; Squire et al. 2004; Luther et al. 2011; Rybakova et al. 2011; Pfuhl et al. 2012). Alternatively, a similar role of parallel elasticity could be played by an I-band spring, like the gigantic protein titin that spans the whole half-sarcomere, connecting the Z line at the end of the sarcomere with the tip of the thick filament and running bound to the surface of the thick filament up to the M-line at the center of the sarcomere (Fig. 1; Maruyama et al. 1977; Wang et al. 1979; Fürst et al. 1988; Linke et al. 2002; Granzier and Labeit 2004). Titin, as an I-band spring, is the only element able to transmit the stress to thick filament also in the resting sarcomere, when no motors are attached to actin, which explains the passive force developed by a resting sarcomere when it is stretched. In frog skeletal muscle, this happens for sarcomere length (SL) > 2.50 μm and reaches ca 0.7 *T*_0_ for SL ~ 3.4 μm (Reconditi et al. 2014). In this respect it must be noted that the cord compliance that can be calculated from the passive force – SL relation is (0.45 μm/(0.7·150 kPa) =) 4300 nm/MPa per hs, ~20 times larger than *C*_p_ determined at full overlap (Fusi et al. 2010, 2017) and even larger than that determined at SL > 2.5 μm (Powers et al. 2017). Such a large difference in the titin-like I band spring finds a molecular explanation in the *in vitro* mechanical studies that allowed the definition of the load dependent structural dynamics of titin (Rivas-Pardo et al. 2016) demonstrating that titin stiffness is modulated in time by the load-dependent equilibrium between folding-unfolding of its immunoglobulin (Ig) domains. This finding suggests that the role of titin as I-band spring in parallel to the array of motors can be much more relevant than that of an elastic element that adds its contribution of force in the extreme condition of a weak half-sarcomere which has undergone a large stretch (Rassier et al. 2005, 2015; Cornachione et al. 2016). Rather titin may work as a dynamic spring that provides a substantial contribution of force to prevent a weak half-sarcomere to give during contraction.

If the contribution of the parallel elasticity to *C*_hs_ were due to either weakly bound motors or any other A-band inter-filament link, the stiffness of the active muscle should reduce at long SL with the reduction of filament overlap. A more complex behavior is expected in the case of a titin-like I-band spring, as in this case the stiffness of the spring may vary with the large changes in its length accompanying the changes in SL. In this respect it is worth to note that, using large stretches, Bagni and co-workers (Bagni et al. 2002) identified an elastic element in parallel with the cross-bridges, defined as a ‘static stiffness’, which rises abruptly upon activation independent of motor attachment and increases with the increase of sarcomere length up to 2.8 μm. Even if the large stretch is a somewhat less direct mean for estimating stiffness changes, as the attached motors are brought into a regime which may imply also detachment-attachment kinetics, the results are intriguing and suggestive of a role of titin in active contraction. A further support to this idea derives from the finding that in vitro titin stiffness increases in the presence of Ca^2+^ (Labeit et al. 2003).

The evidence, in both resting and active muscle, for the presence of an I-band spring element like titin that anchors the Z line to the end of the thick filament, makes FHS 1981 model no longer valid, because it assumes that, at any SL, the absence of cross-bridges results in an infinite half sarcomere compliance. The attempts to consider the effect of titin on the dynamics of the half-sarcomere done so far have used the simplified assumption that it is in parallel with the force generating cross-bridges (Rice et al. 2008; Campbell et al. 2018). However, an I-band spring as titin is neither in parallel (sharing the same length change) nor in series (sharing the same force change) with any other element in the hs.

Here we integrate the FHS1981 model to include for the first time the contribution of an I-band spring to the half sarcomere compliance in its proper configuration. The model represents a tool for the interpretation of measurements of hs stiffness at increasing SL, which is important either in relation to the mechanism of stabilisation of SL against the consequence of sarcomere inhomogeneity in active force generation, or for investigations on the role of titin as mechano-sensor in thick filament regulation (Linari et al. 2015; Reconditi et al. 2017; Piazzesi et al. 2018).

## METHODS

In the FHS 1981 model, assuming that the compliance of the array of cross-bridges is not smaller than the cumulative compliance of the actin and myosin filaments, *C*_hs_ can be calculated by the sum of the equivalent compliances of the three elements:

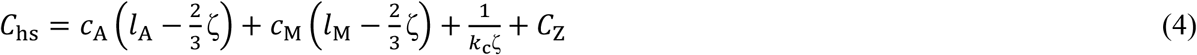

Where *c*_A_ and *c*_M_ are the compliances per unit length of the thin and thick filaments respectively, *k*_c_ is the stiffness per unit length of the array of the cross-bridges, *C*_Z_ is the compliance of the Z line, *l*_A_ and *l*_M_ are the length of the thin and thick filament respectively, ζ is the length of overlap of thin and thick filaments in each half-sarcomere and 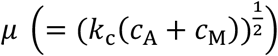 is a parameters that increases as the cumulative compliance of the filaments increases relative to that of the motor array (1/*k*_c_).

Provided that *μ*ζ/2 is not too large, Eq. 1 simplifies to:

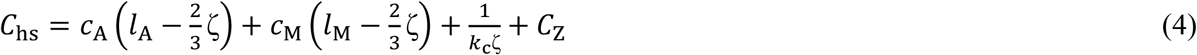

Given the small axial extension of the Z line (~30–50 nm in the fast skeletal muscles; Luther 2009), its contribution *C*_Z_ to *C*_hs_ will be neglected hereafter. Estimates of *c*_A_, *c*_M_ and *k*_c_ obtained from both X-ray diffraction and mechanical measurements (Huxley et al. 1994; Wakabayashi et al. 1994; Reconditi et al. 2004; Huxley et al. 2006; Piazzesi et al. 2007; Fusi et al. 2014; Brunello et al. 2009) indicate that the condition *μ*ζ/2 <1 is fulfilled. Consequently, according to the model, *C*_hs_ can be considered functionally as the series of two compliances: the myofilament compliance, 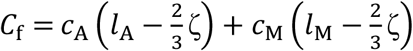 and the motor compliance, 1/*k*_c_ζ.

According to the model of FHS1981 (Fig. 2), an external force *T* applied to the hs is entirely born by the thick and thin filaments in the regions that do not overlap and is shared between the two filaments in the overlap region, where the cross-bridges transfer force between the two filaments. At any axial position the force born by the thin filament, *T*_A_, and that born by the thick filament, *T*_M_, add to give the total force *T* applied to the hs. Taking ξ as the axial distance measured from the center of the overlap zone, and positive toward the Z line, the difference d*T*_A_(ξ)= *T*_A_(ξ+d ξ)-*T*_A_(ξ) in the force born by the thin filament along a small axial distance dξ at the coordinate ξ is *k*_c_ dξ *x*(ξ), where *k*_c_ dξ is the stiffness of the small cross-bridge segment dξ and *x*(ξ) is the distortion of the cross-bridge array at that coordinate, taken positive for stretches (Fig. 3, upper panel). In the same way, for the thick filament d*T*_M_(ξ) = -*k*_c_dξ*x*(ξ), consistent with the fact that force is transferred between the two filaments through the cross-bridge array and, in each point along the hs, is the same and shared between the filaments in the overlap region. This leads to Eq. A1 of FHS1981:

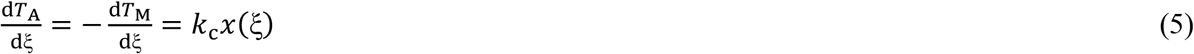

**Fig. 2.**
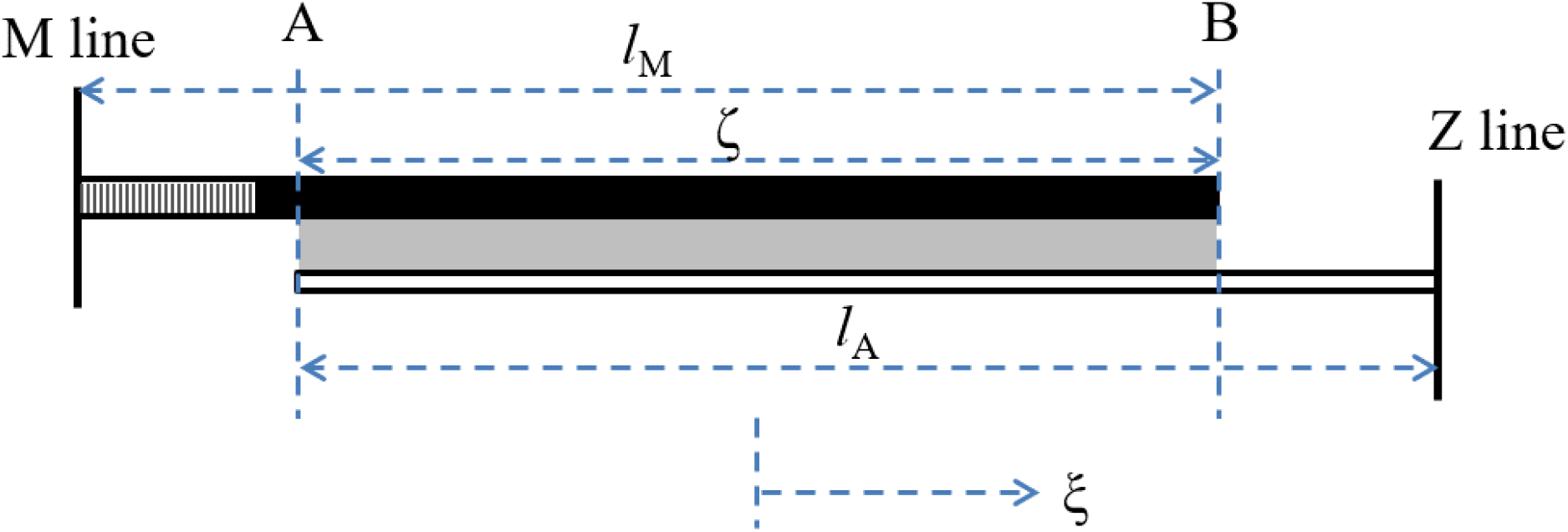
Diagram indicating the arrangement of the thick and thin filaments and the cross-bridges in the half-sarcomere. The half-thick filament (length *l*_M_; black, where myosin motors are present; dashed, bare zone, where no myosin motors are present) extends from the M line, at the center of the sarcomere. The grey band represents the cross-bridges in the region (length ζ) that overlaps with the thin filament (white; length *l*_A_) that extends from the Z line. Motors attaching to the thin filament (cross-bridges) contribute to the compliance of the half-sarcomere. ξ is the axial coordinate, with the origin in the middle of the overlap region (adapted from Ford et al. 1981).

**Fig. 3.**
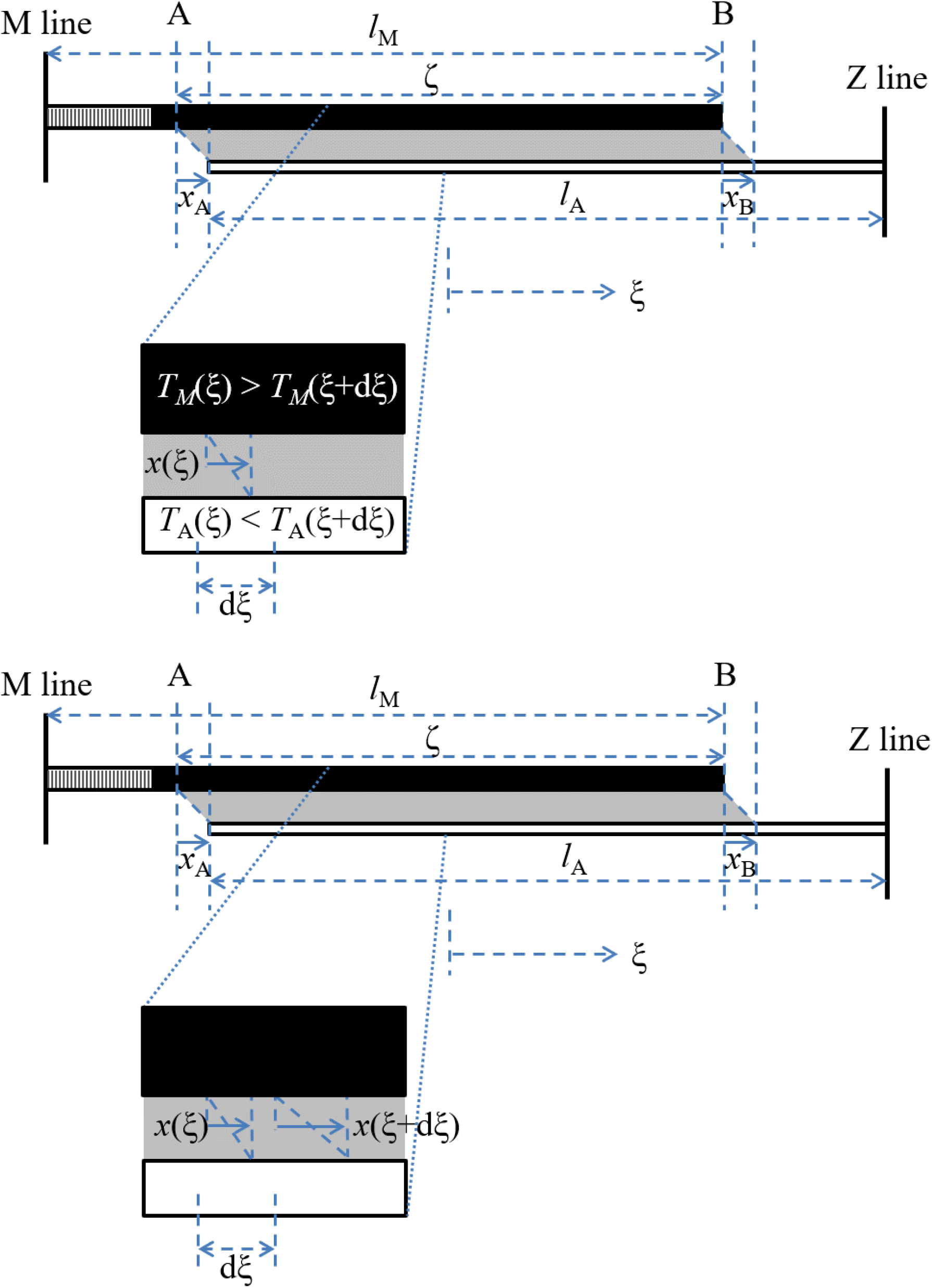
Schematic of the half sarcomere strained by an external force *T*. *x*_A_ and *x*_B_ are the cross-bridge distortion at the point A and B along the half-sarcomeres, at coordinate ξ = -ζ/2 and ξ = +ζ/2 respectively. Upper panel: the expanded region around the generic coordinate ξ shows that the force in a small region dξ is transmitted between the thick and thin filament through a distortion *x*(ξ) of the cross-bridge array segment dξ wide and *k*_c_dξ stiff. Lower panel: in the expanded region is now indicated the different distortion *x* of the cross-bridge array over a small axial distance dξ.

Over a small axial distance dξ, the distortion of the cross-bridges differs by a quantity d*x*(ξ) = *x*(ξ + dξ) − *x*(ξ) that results from the different strain in the thin and thick filaments (Fig. 3, lower panel), so that:

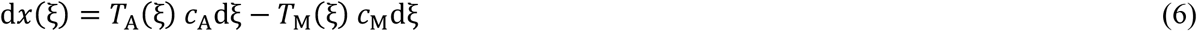

and thus:

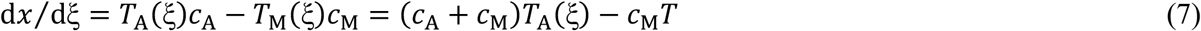

that is Eq. A2 of FHS1981.

Differentiating (7) and substituting for d*T*_A_/dξ from Eq. 5 gives

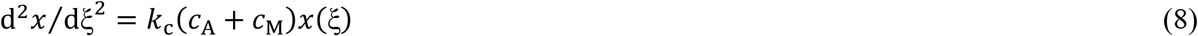

whose solution (Eq. A4a of FHS81) is

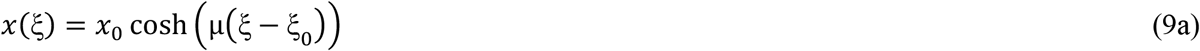

where

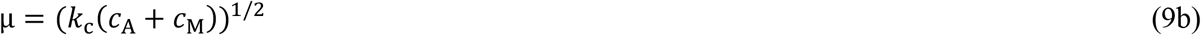

The presence of an elastic link between the end of the thick filament and the Z line, which is the role played by titin in the I band, changes the boundary conditions with respect to FHS1981 model, and in turns affects the expressions for *x*_0_ and ξ_0_. In the original model, at point A (Fig. 2) 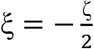 and 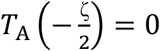 and at point B 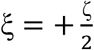 and 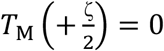.

In the presence of titin link, the force on the thick filament at B, i.e. at the end of the filament, is the same as that, *T*_T_, born by titin in the I band. This force, added to *T*_A_ in the I band, gives the full force *T* found at the Z line:

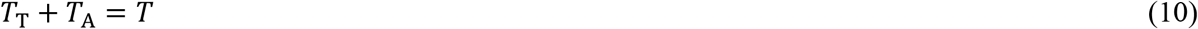

Thus, to determine the boundary condition at *B*, one must first determine how *T* is distributed between thin filament and titin in the I band. To do that, we consider that under a force *T* on the hs, in the I band the strain of titin, *c*_T_(*l*_A_ − ζ)*T*_T_, where *c*_T_ indicates the compliance of titin per unit length, must be the same as the strain of the thin filament, *c*_A_(*l*_A_ − ζ)*T*_A_, plus the strain of the cross-bridges at *B*, *x*_B_ (Fig. 4):

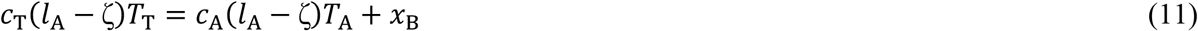

**Fig. 4.**
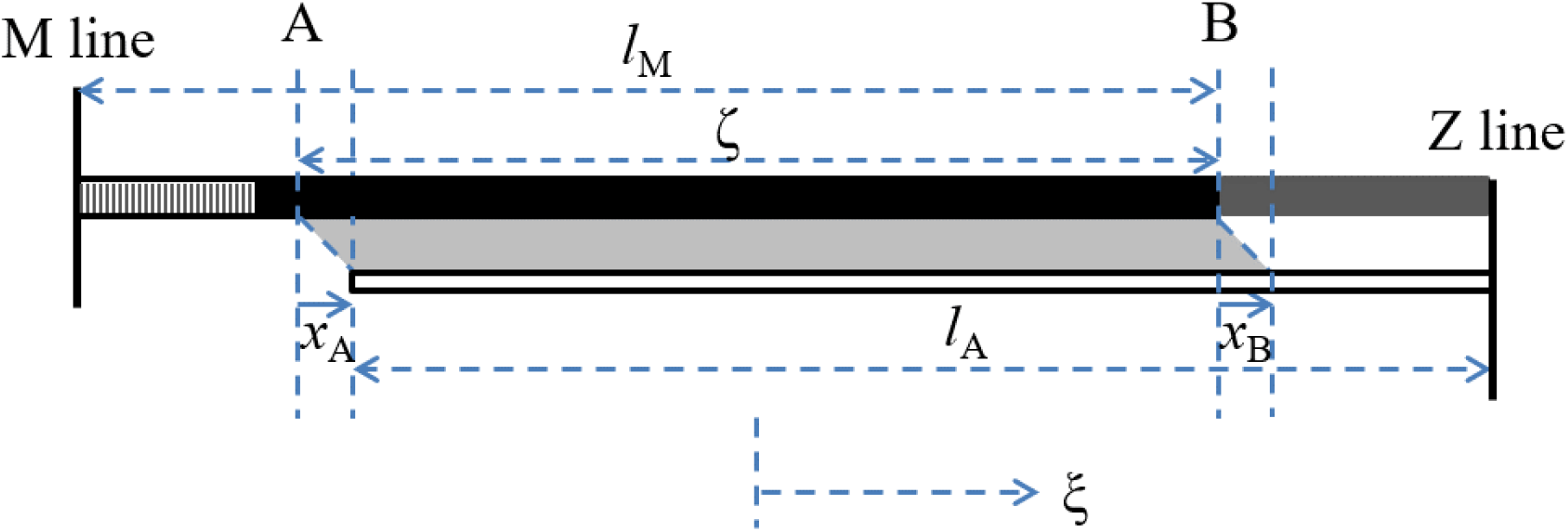
Diagram of the half-sarcomere incorporating the titin (dark gray band, extending from the Z line to the edge of the thick filament). When an axial force is applied to the half-sarcomere, the strain of the titin is equal to the strain of the thin filament in the I band plus the strain of the cross-bridge array at B, *x*_B_.

Eq. 10 and Eq. 11 provide the expression for *T*_A_ and *T*_T_ in the I band (and thus their values in B) as:

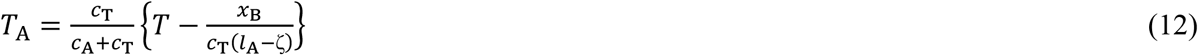

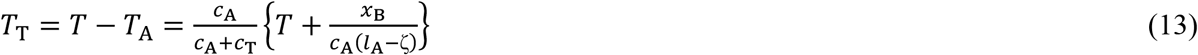

With this, the boundary conditions in A and B become respectively:

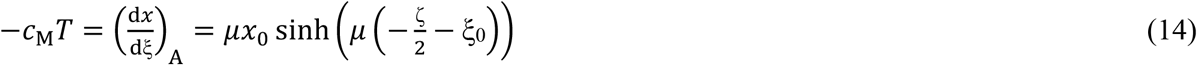

and

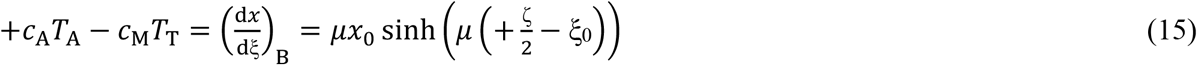

By using Eq. 11 and Eq. 12, Eq. 15 can be rearranged to:

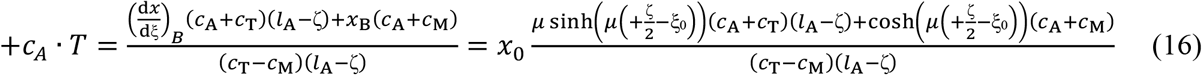

Eq. 14 and Eq. 16 are then equivalent to Eq. A5a and Eq. A5b of FHS1981 respectively.

Equating Eq. 14 and Eq. 16 for *T* leads to:

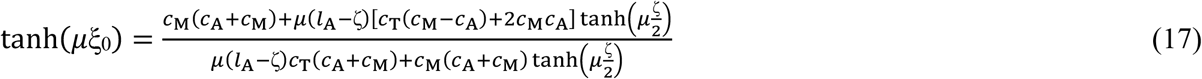

Subtracting Eq. 14 from Eq. 16 gives:

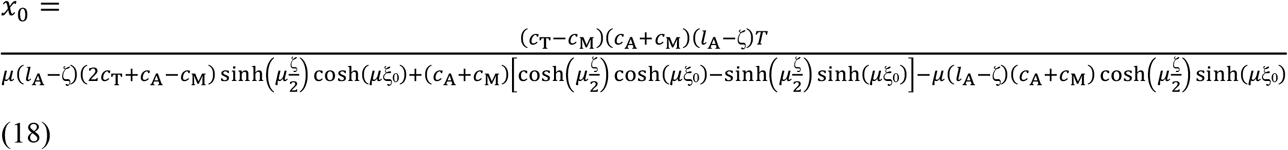

Eq. 17 and Eq. 18 are equivalent to Eq. A6 and Eq. A7 of FHS1981 respectively.

The compliance per half sarcomere, *C*_hs_, is obtained by calculating the total strain *S*_hs_ that an external force *T* induces on the half-sarcomere, and dividing the result by *T*, according to the relation *C*_hs_= *S*_hs_/*T.*

Going from the M line to the Z line there are four possible paths to calculate *S*_hs_, and of course the results are the same.

The same path as in FHS1981 (where the possible paths are just two, since there is no titin link) is followed here. *S*_hs_ is given by the sum of the strain of the thick filament within the H zone (from the M line to the beginning of the overlap with the thin filament), the displacement of tip of the thin filament with respect to thick filament (or the strain of cross-bridges array at A, *x*_A_), the extension of the thin filament in the overlap region, and the extension of thin filament within the I-band (Fig. 4):

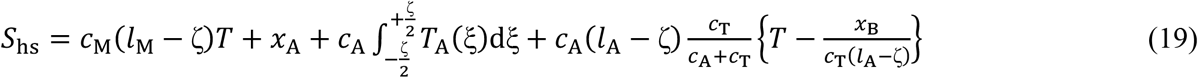

Eq. 15 is equivalent to Eq. A8 in FHS1981, and similarly, from Eq. 7:

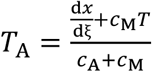

and the integral in Eq. 19 becomes:

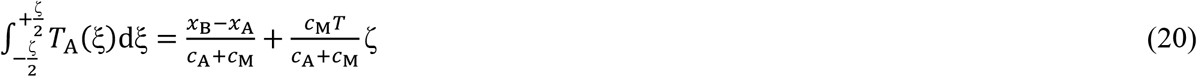

Thus:

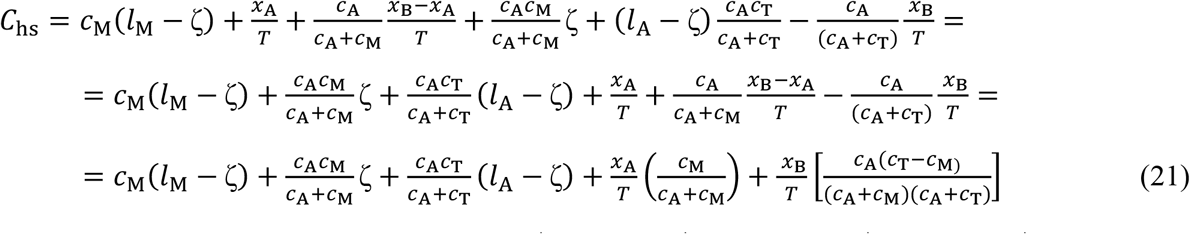

Finally, substituting *x*_A_ and *x*_B_ with 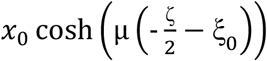 and 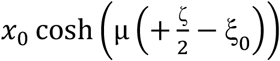 respectively, and using Eq. 17 and Eq. 18 lead to:

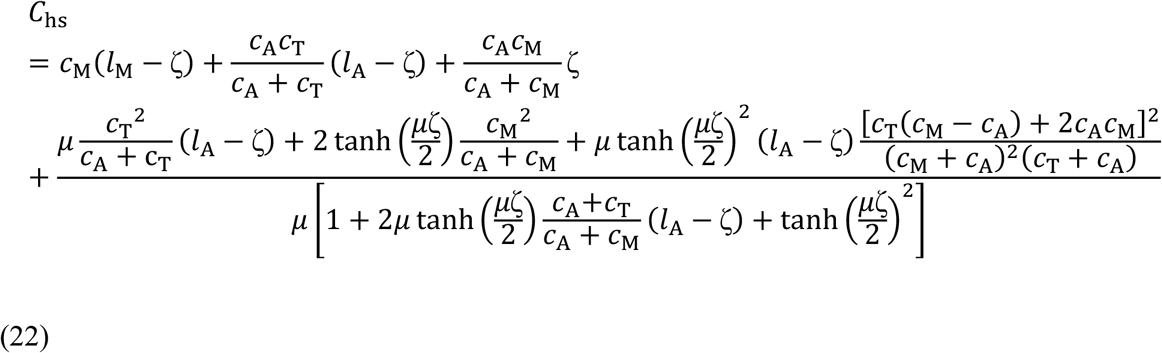

Provided that *μ*ζ/2 is not too large, dividing both numerator and denominator of the last term for tanh(*μ*ζ/2) and using the approximations:

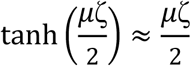

and

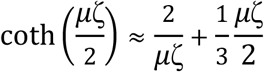

Eq. 22 can be approximated by:

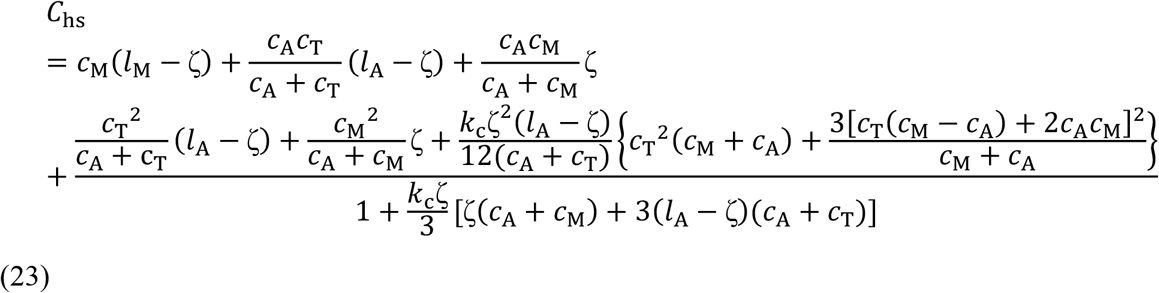

where the dependence of *C*_hs_ on *k*_c_, the stiffness per unit length of the motor array, is made explicit.

## RESULTS

When *c*_T_ → ∞, Eq. 22 and Eq. 23 reduce to Eq. A9 and Eq. A10 of FHS1981, as expected. On the contrary, when *k*_c_, and then μ, → 0 or *c*_A_ → ∞, while FHS1981 predicts *C*_hs_ → ∞, Eq. 22 predicts *C*_hs_ = *c*_M_*l*_M_ + *c*_T_(*l*_A_ − ζ), i.e. the series of thick filament and I-band titin compliances. These results are consistent with the structural/mechanical models drawn in Fig. 2 and Fig. 4 respectively.

The main difference introduced by the I-band spring that links the edge of the thick filament to the Z line is that in this case the compliance of the half-sarcomere cannot any longer be thought as the sum of compliances in series, even within the approximation that leads to Eq. A10 of FHS1981. Thus, while FHS1981 allowed to define an equivalent filament compliance *C*_f_ contributing to *C*_hs_, as represented by Eq. 3, when the contribution of titin is considered, the concept of equivalent filament compliance is no longer applicable.

Experimentally, the values of the parameters *c*_A_, *c*_M_, *c*_T_, *k*_c_ can be estimated, for example, by applying fast (4 kHz) length oscillations to a muscle fiber during the rise of an isometric tetanus (Fusi et al. 2017) to measure *C*_hs_ at several force levels during the rise of an isometric tetanus. The number of attached motors (*n*_A_) varies linearly with force (Piazzesi et al. 2018), and so does their stiffness: *k*_c_ζ=*k*_c0_ζ*T*, where *k*_c0_ is the stiffness per unit length of the cross-bridge array at the maximal isometric force *T*_0_, and *T* is the force during the tetanus rise expressed in *T*_0_ units. In this way, the values of *C*_hs_ at different *T* can be fitted by Eq. 22 or Eq. 23 to determine the different parameters.

As an example, to evaluate the possible contribution of titin to the half-sarcomere compliance, we have calculated how *C*_hs_ from Eq. 22 should vary during the tetanus rise with the values for *c*_A_, *c*_M_ and *k*_c0_ as estimated in Brunello et al. (2014), and with different values for *c*_T_. Here we neglect the possible contribution of the compliance in parallel with the cross-bridges (*C*_p_). In Brunello et al. (2014) the values of the myofilament compliances are estimated by means of X-ray diffraction measurements as *c*_A_ = 2.29 nm/*T*_0_/μm and *c*_M_ = 2.80 nm/*T*_0_/μm, and, by comparing these results with mechanical measurements, *k*_c0_ is calculated to be 0.806 *T*_0_/nm/μm. The results are reported in Fig. 5, where *C*_hs_ as a function of *T* is plotted for different *c*_T_ (1, 10, 100 and 1000 nm/*T*_0_/μm) at four different sarcomere lengths (SL), 2.15 (A), 2.5 (B), 2.7 (C) and 2.90 μm (D) (and correspondingly different ζ from 0.7 to 0.325 μm, having taken *l*_A_=0.975 μm and *l*_M_=0.8; Brunello et al. 2014). It can be seen that the presence of a titin-like compliance with *c*_T_ of 10 nm/*T*_0_/μm or less (short-dashed lines) is not compatible with the observed *C*_hs_-*T* relation at 2.15 μm (Fusi et al. 2014), as in this case, unlike what observed, *C*_hs_ at lower forces is systematically reduced with respect to the value in the absence of the titin-like spring to a value that is almost constant independent of isometric force, which is directly proportional to *n*_A_. Instead for higher *c*_T_ values (100 nm/*T*_0_/μm or larger, long-dashed lines), *C*_hs_ at high forces (or *n*_A_) is unaffected, while at low forces it is reduced by an extent that is larger at smaller forces in agreement with experimental results at 2.15 um SL (11) and, at any force, is larger at longer SL. Superimposed *C*_hs_-force relations at the four SL are shown in Fig. 6 in the absence of a titin-like spring (A, B) and with a titin-like spring with *c*_T_ 100 nm/*T*_0_/μm (C, D). In the expanded scale it can be seen that in the absence of titin (B) the increase in *C*_hs_ with SL (that is with the reduction of filament overlap) present at high forces becomes progressively smaller with the reduction of force, since *C*_hs_ tends to ∞ as *n*_A_ tends to zero. Instead, in the presence of a titin-like spring with *c*_T_ 100 nm/*T*_0_/μm, the SL-dependent upward shift in *C*_hs_ remains at any force, as at low force the contribution of titin takes over that of *n*_A_.

**Fig. 5.**
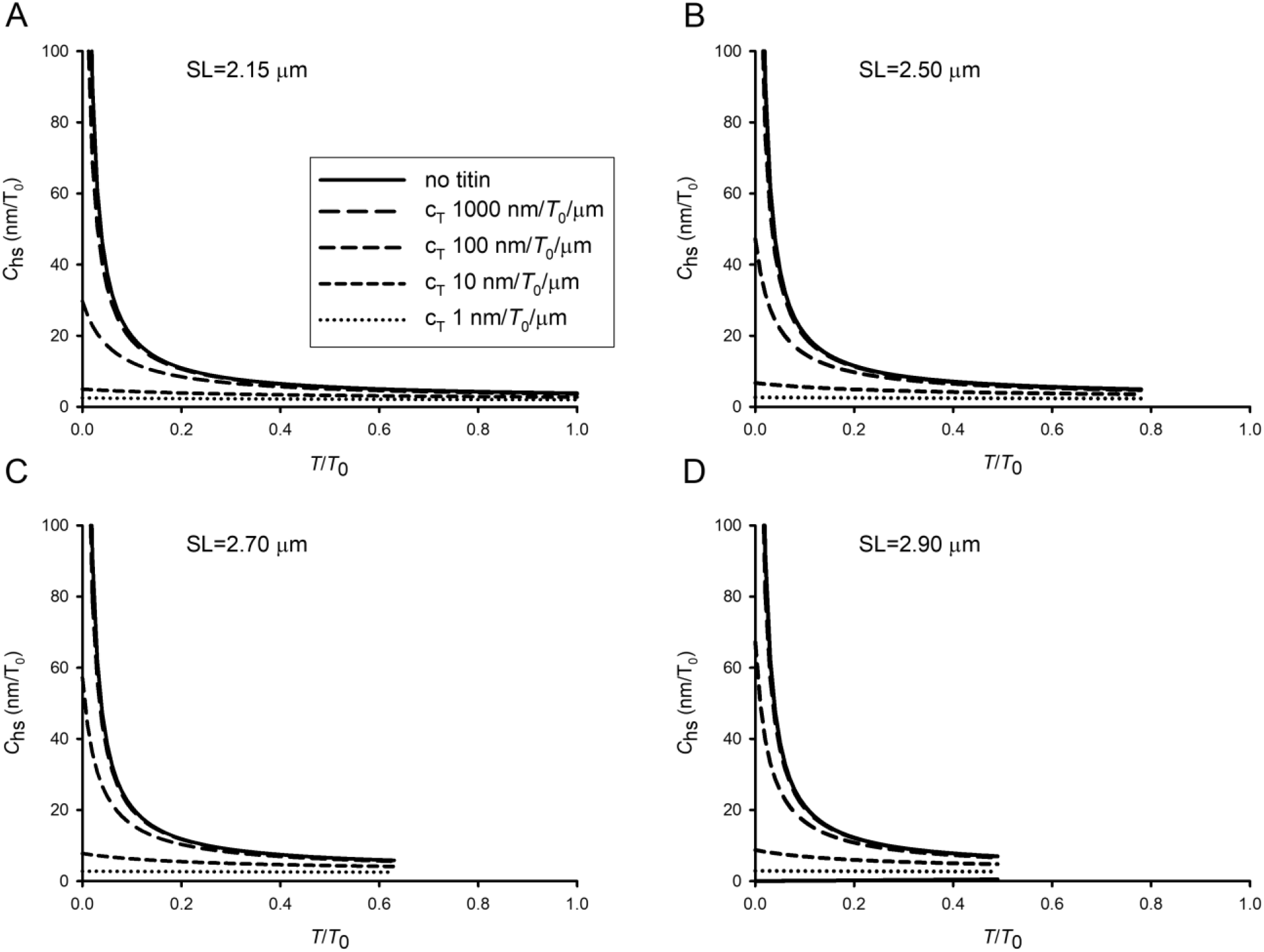
Compliance of the half sarcomere *C*_hs_ calculated as a function of force *T* during the rise of an isometric tetanus for different values of titin compliance per unit length (*c*_T_, as in the inset) and at four different sarcomere lengths (A. SL = 2.15 μm; B. SL = 2.50 μm; C. SL = 2.70 μm; D. SL = 2.90 μm). *T*_0_ is the maximal tetanic force developed in isometric contraction at SL = 2.15 μm. *c*_A_, *c*_M_ and *k*_c_ from Brunello et al. 2014 (see text). *l*_A_ = 0.975 μm, *l*_M_ = 0.8 μm.

**Fig. 6.**
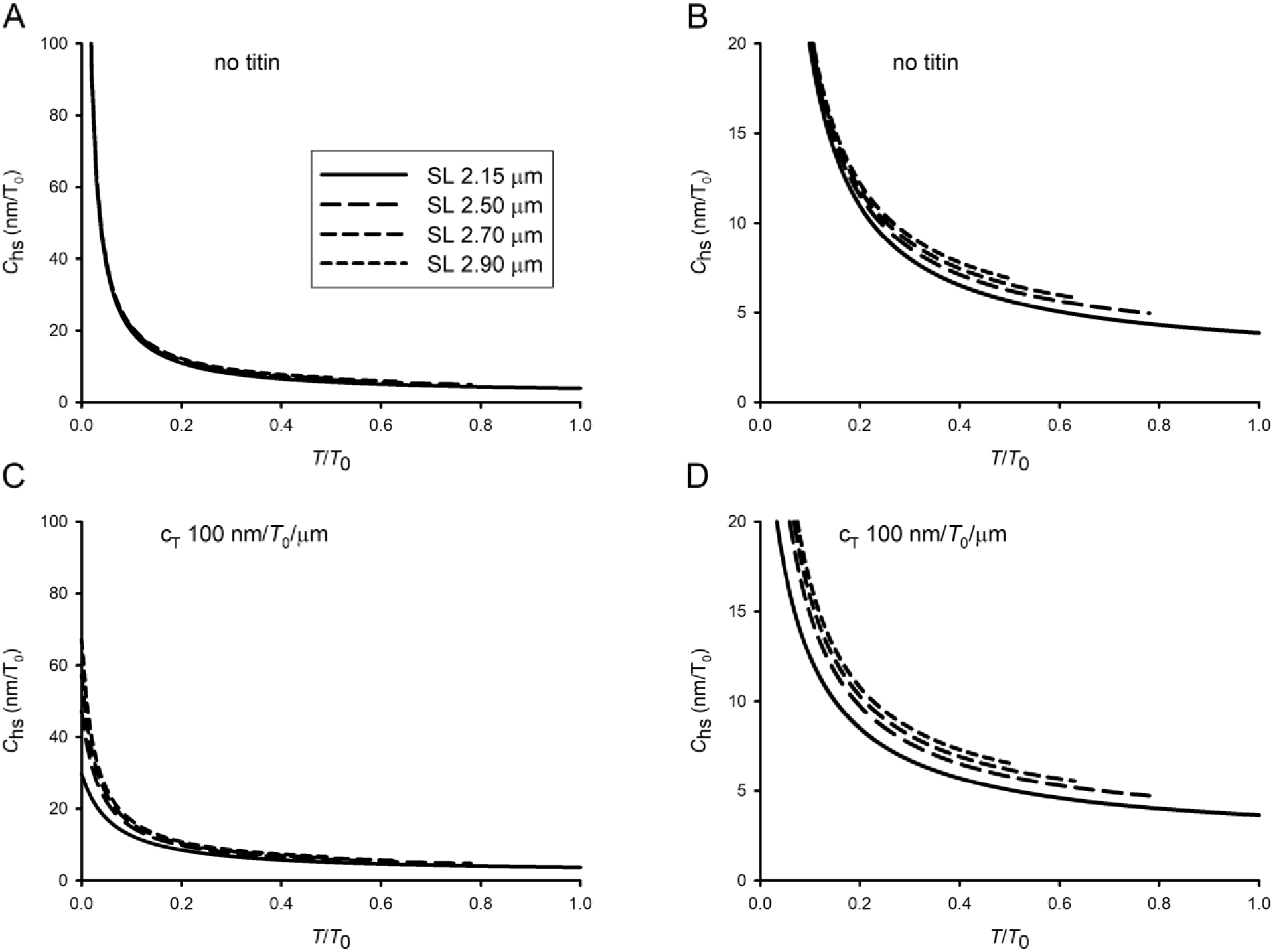
Compliance of the half-sarcomere *C*_hs_ calculated as a function of force *T* during the rise of an isometric tetanus at different sarcomere lengths (SL, as in the inset) in the absence (A, B) or in the presence (C, D) of titin with a compliance per unit length *c*_T_ = 100 nm/*T*_0_/μm.

## DISCUSSION

The contribution to the compliance of a titin-like I-band spring emerges whenever, like at low level of force during the rise of an isometric tetanus, the number of attached motors *n*_A_ is relatively small with respect to those attached at the isometric tetanus plateau. In this way the I-band spring prevents *C*_hs_ to rise to infinite as *n*_A_ approaches zero (Fig. 5). The reduction of Chs is larger if the compliance of the titin-like element (*c*_T_) is smaller and, for a given value of *c*_T_, is larger at lower forces. This is emphasised in Fig. 7A, where the points (extracted by the relations in Fig. 5) indicate the relation between *C*_hs_ and *c*_T_ at 0.5 *T*_0_ (triangles) and 0.1 *T*_0_ (circles). The reduction in *C*_hs_ produced by introducing a titin-like spring with a *c*_T_ of 100 nm/*T*_0_/μm is estimated at 0.1 *T*_0_ and at 0.5 *T*_0_ by the length of the segment *m* and *n* respectively, showing that the effect of titin is 12 times larger at 0.1 than at 0.5 *T*_0_. In this respect the contribution of an I-band spring like titin is similar to that of an A-band spring as that represented by a fraction of no-force generating or weakly-bound motors in parallel with the array of force generating motors (Colombini et al. 2010; Fusi et al. 2017).

**Fig. 7.**
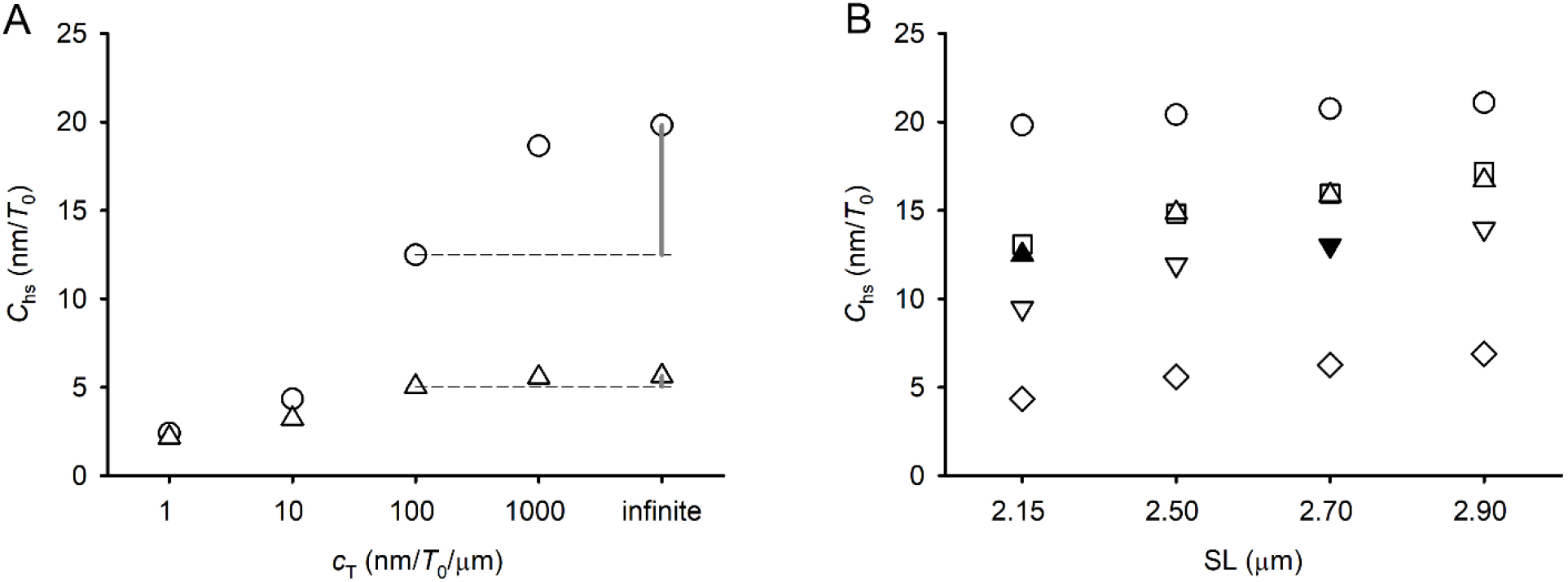
A. Compliance of the half-sarcomere *C*_hs_ calculated as a function of titin compliance per unit length, *c*_T_, at SL 2.15 μm at forces 0.1 *T*_0_ (circles) and 0.5 *T*_0_ (triangles) during the rise of an isometric tetanus. The vertical gray lines indicate the reduction in *C*_hs_ caused by introducing an I-band spring with *c*_T_ = 100 nm/*T*_0_/μm at either 0.1 *T*_0_ (m) or 0.5 *T*_0_ (n). B. Compliance of the half-sarcomere *C*_hs_ calculated as a function of SL at 0.1 *T*_0_. Circles: *C*_hs_ contributed only by the myofilaments and cross-bridges; squares: effect of a parallel elastic element with compliance *C*_p_ as described in the text; triangles, reverse triangles and diamonds: effect of titin-like I-band spring with compliance *c*_T_ = 100 nm/*T*_0_/μm, 50 nm/*T*_0_/μm and 10 nm/*T*_0_/μm respectively. See text for filled symbols.

The effect on *C*_hs_ of an I-band spring like titin with respect to that of an A-band spring may become quite more specific in relation to changes in sarcomere length. The question is analysed in Fig. 7B by comparing the relations between *C*_hs_ and SL at force 0.1 *T*_0_, at which both an I-band and an A-band spring produce a large effect, obtained either with the contribution of cross-bridges in the absence of any added spring (circles) or in the presence of either an A-band spring like that determined in Fusi et al. (2014) (squares, *C*_p_ = 29 nm/*T*_0_) or an I-band spring with *c*_T_ of 100 nm/*T*_0_/μm (triangles), 50 nm/*T*_0_/μm (reverse triangles) and 10 nm/*T*_0_/μm (diamonds). At SL 2.15 μm an A-band spring with *C*_p_ 29 nm/*T*_0_ decreases *C*_hs_ by almost the same amount (~35%) as an I-band spring with *c*_T_ 100 nm/*T*_0_/μm. In fact, taking into account the length of the I-band spring (SL/2 – *l*_M_ = 0.275 μm), the actual I-band spring compliance *C*_T_ is (100 nm/*T*_0_/μm·0.275 μm =) 27 nm/*T*_0_, not significantly different from *C*_p_ of the A-band spring determined in Fusi et al. (2014) at SL 2.15 μm (square). Thus, both the I-band and the A-band springs with constant stiffness do contribute to increase *C*_hs_ with the increase in SL, as demonstrated by the finding that the slopes of both *C*_hs_-SL relations identified by squares and triangles are larger than the slope of the *C*_hs_ – SL relation calculated with the contribution of cross-bridges without any added spring (circles). The *C*_hs_-SL relation is shifted progressively downward with the reduction of *c*_T_ to 50 nm/*T*_0_/μm (reverse triangles) and 10 nm/*T*_0_/μm (diamonds).

It must be noted that the relations in Fig. 7B represent the theoretical predictions of the effect on *C*_hs_ of springs that have a constant compliance per unit length. To apply these predictions to the contribution to *C*_hs_ of a titin-like I-band spring in situ, we may need to take into account of experimental evidence (first of all the passive force-SL relation, but also the response of active muscle to sudden larger stretch (Bagni et al. 2002)) showing that titin stiffness increases at larger SL. Actually, yet neither in situ nor in vitro measurements of SL dependence of titin stiffness have been performed with small length perturbation in the ≥ 4kHz frequency domain, which should prevent the confounding effects of the faster components of the relaxation processes related to titin unfolding (Kellermayer et al. 1997; Rivas-Pardo et al. 2016) and motor detachment/attachment (Lombardi and Piazzesi 1990). It looks likely, anyway, that also in the active muscle, as in the resting muscle, titin stiffness increases with SL. Under these conditions the relations in Fig. 7B provide a fundamental tool to interpret the changes in *C*_hs_ related to the contribution of a titin-like I-band spring. As an example, let’s assume that *c*_T_ at SL 2.15 μm is 100 nm/*T*_0_/μm and compare the *C*_hs_ attained at SL 2.7 μm with *c*_T_ 100 nm/*T*_0_/μm (triangle) to that with *c*_T_ 50 nm/*T*_0_/μm (reverse triangle). If *c*_T_ does not change with the increase in SL, *C*_hs_ at 2.7 μm (triangle) increases by 25%, as the result of both the reduction of *n*_A_ (due to the reduction of filament overlap by 0.25 nm) and doubling of the length of the I-band spring, (SL/2 – *l*_M_), from 0.275 to 0.55 μm. However, if we assume that titin is a tunable spring which perfectly adapts its stiffness to the large changes in the I-band length, so that its *c*_T_ is halved when its length is doubled, then by increasing the SL from 2.15 to 2.7 μm (which doubles the I-band spring length) the change in *C*_hs_ would be quite smaller as it would depend only on the reduction in number of attached motors with reduction in overlap, as shown by comparing the ordinate value of the filled triangle (*c*_T_ 100 nm/*T*_0_/μm, SL 2.15 μm) with the filled reverse triangle (*c*_T_ 50 nm/*T*_0_/μm, SL 2.7 μm).

## ACKNOWLEDGMENT

We thank Vincenzo Lombardi for insightful comments on the manuscript. This work was supported by University of Florence (competitive project rictd1819) (Italy).

## AUTHOR CONTRIBUTIONS

M.R. developed the mathematical formalism. I.P. made the numerical calculations. M.C. and M.R. wrote the manuscript.

